# Antibody-induced crosslinking and cholesterol-sensitive, anomalous diffusion of nicotinic acetylcholine receptors

**DOI:** 10.1101/744664

**Authors:** Alejo Mosqueira, Pablo A. Camino, Francisco J. Barrantes

**Author notes:** Address correspondence to: F.J. Barrantes, Laboratory of Molecular Neurobiology, BIOMED UCA-CONICET, Av. Alicia Moreau de Justo 1600, C1107AFF, Buenos Aires, Argentina. The abbreviations used are: BTX, α-bungarotoxin; mAb, monoclonal antibody; nAChR, nicotinic acetylcholine receptor; CDx, methyl-β-cyclodextrin; CDx-Chol, cholesterol-methyl-β-cyclodextrin complex; MSD, mean-squared displacement; SPT, single-particle tracking.

## Abstract

Synaptic strength depends on the number of cell-surface neurotransmitter receptors in dynamic equilibrium with intracellular pools. Dysregulation of this homeostatic balance occurs e.g. in myasthenia gravis, an autoimmune disease characterized by a decrease in the number of postsynaptic nicotinic acetylcholine receptors (nAChRs). Monoclonal antibody mAb35 mimics this effect. Here we use STORM nanoscopy to characterize the individual and ensemble dynamics of mAb35-crosslinked receptors in the clonal cell line CHO-K1/A5, which robustly expresses adult muscle-type nAChRs. Antibody labeling of live cells results in 80% receptor immobilization. The remaining mobile fraction exhibits a heterogeneous combination of Brownian and anomalous diffusion. Single-molecule trajectories exhibit a two-state switching behavior between free Brownian walks and anticorrelated walks within confinement areas. The latter act as permeable fences (∼34 nm radius, ∼400 ms lifetime). Dynamic clustering, trapping and immobilization also occur in larger nanocluster zones (120-180 nm radius) with longer lifetimes (11 ± 1 s), in a strongly cholesterol-sensitive manner. Cholesterol depletion increases the size and average duration of the clustering phenomenon; cholesterol enrichment has the opposite effect. The disclosed high proportion of mAb35-crosslinked immobile receptors, together with their anomalous, cholesterol-sensitive diffusion and clustering, provides new insights into the antibody-enhanced antigenic modulation that leads to physiopathological internalization and degradation of receptors in myasthenia.

A preliminary version of this work has appeared in the biorXiv repository: https://www.biorxiv.org/content/10.1101/744664v1. The study was not pre-registered.

## Introduction

The nicotinic acetylcholine receptor (nAChR) is one of the most ubiquitous neurotransmitter receptor proteins in both peripheral and central nervous systems of vertebrates (for a recent review see (Changeux 2018)). This pentameric ligand-gated ion channel intervenes in a variety of physiological processes and is subject to dysfunction in pathological conditions such as chronic pain, schizophrenia spectrum disorders, Alzheimer disease, Parkinson disease, or neurological autoimmune diseases in the case of brain disorders. In the peripheral nervous system, the nAChR is also the target of autoimmune diseases like myasthenia gravis, an invalidating disease of the neuromuscular junction (NMJ), the peripheral cholinergic synapse of vertebrates (Vincent 2002; Paz and Barrantes 2019). In the latter, about two-thirds of the autoantibodies found in myasthenic patients are directed against the so-called main immunogenic region (MIR) of the muscle-type nAChR (Tzartos et al. 1988). This finding resulted from the introduction of libraries of cloned hybridoma cell lines producing monoclonal antibodies to individual nAChR subunits and regions thereof (Tzartos and Lindstrom 1980). mAb35, the antibody used in the present study, is an IgG1 immunoglobulin isotype reacting with the MIR epitope located on the extracellular surface of the nAChR α1 subunit of the muscle-type nAChR (Tzartos et al. 1991). Remarkably, this monoclonal antibody was originally raised against the nAChR of Electrophorus electricus, the elasmobranch electric eel, and yet it competes with >75% of the serum autoantibodies obtained from humans suffering from myasthenia gravis (Tzartos et al. 1982). MIR epitopes recognized by rat mAbs consist of two discontiguous sequences, which are adjacent in the native conformation and confer high-affinity binding for rat monoclonal antibodies (Luo et al. 2009). A single IgG monoclonal antibody molecule can crosslink two nAChR molecules (Tzartos and Starzinski-Powitz 1986). Recent X-ray structural studies have confirmed that a single mAb35 molecule is able to bind two α1 subunits from two adjacent monomers-thereby crosslinking them-whereas it does not bind to the two α1 subunits within a single nAChR monomer (Noridomi et al. 2017). Moreover, antibodies directed against the MIR of the receptor have been reported to form oligomeric complexes larger than the dimer (Conti-Tronconi et al. 1981; Tzartos et al. 1986).

Ultracentrifugation and transmission electron microscopy (Barrantes 1982) and atomic force microscopy (Pissinis et al. 2015) techniques have made apparent the wide range of oligomeric forms that the nAChR can adopt in non-ionic detergent, ranging from monomers to dimers and higher oligomeric species. This structural repertoire possibly mimics a similar range of oligomeric states of the protein in situ, given the high-packing densities of the receptor in the native post-synaptic membrane, maximizing in-plane receptor-receptor contacts that favor oligomerization.

Here, we set out to explore the effects of mAb35-mediated crosslinking of the adult muscle-type nAChR heterologously expressed in a mammalian cell system on the translational dynamics and spatial organization of the protein imaged in live CHO-K1/A5 cells using STORM (stochastic optical reconstruction microscopy), single-molecule tracking and a combination of analytical and statistical techniques. We also analyze the impact of acute membrane cholesterol depletion with methyl-β-cyclodextrin (CDx) or enrichment (CDx-cholesterol complexes, CDx-Chol) on the diffusion of the receptor. We account for the heterogeneous, cholesterol-dependent diffusion behavior of the nAChR in physical terms by a two-state switching model (Grebenkov 2019) between free Brownian walks and obstructed diffusion within confinement sojourns. Thus, the disclosed anomalous diffusion of the receptor crosslinked by a subclass of IgG1 antibodies known to produce severe loss of nAChR by complement-mediated lysis of the postsynaptic membrane (Vincent 2002; Paz and Barrantes 2019), appears as one of the earliest links in the chain of physiopathological events responsible for antigenic modulation, the cascade leading to enhanced nAChR internalization and degradation, and complement-mediated lysis of the postsynaptic membrane observed in myasthenia.

## Materials and Methods

Mouse monoclonal antibody clone mAb35 (purified immunoglobulin, product No. M-217) against the extracellular moiety of the nicotinic acetylcholine receptor (nAChR) α_1_-subunit, methyl-β-cyclodextrin (CDx, product No. C4805), catalase from Aspergillus niger (product No. C3515), glucose oxidase Type VII (product No. G2133), β-mercaptoethanol (product No. 63689), and polyvinylalcohol (PVA, 25,000 MW, prod. No. 184632) were purchased from Sigma Chem. Co. (St. Louis, MO). Texas Red-labeled anti-IgG (product. No. T-862) secondary antibodies were purchased from ThermoFisher Sci., Invitrogen Argentina.

## Methods

### Cell culture

CHO-K1/A5 cells, a clonal cell line robustly expressing adult muscle-type nicotinic acetylcholine receptor (nAChR) (Roccamo et al. 1999) were grown in Ham’s F12 medium supplemented with 10% fetal bovine serum for 2-3 days at 37°C before experiments. Cells are used for a maximum of 20 passages. No ethical approval was required for the use of the biological material.

### Acute cyclodextrin-mediated cholesterol depletion/enrichment of cultured cells

Acute cholesterol depletion/enrichment was performed prior to fluorescent labeling by treating CHO-K1/A5 cells with 10-15 mM CDx or CDx-cholesterol (CDx-Chol) complexes in Medium 1 (“M1”: 140 mM NaCl, 1 mM CaCl_2_, 1 mM MgCl_2_ and 5 mM KCl in 20 mM HEPES buffer, pH 7.4) (Borroni et al. 2007; Almarza et al. 2014). Samples were taken at 20 min from culture dishes incubated at 37°C in the presence or absence of the cholesterol-modifying chemical.

### Single-molecule localization superresolution microscope setup

The optical nanoscope constructed in our laboratory operates in the stochastic optical reconstruction microscopy (STORM) (Rust et al. 2006) /ground state depletion microscopy followed by individual molecule return (GSDIM) (Folling et al. 2008) modes. A 532 nm pumped solid state DPSS 300 mW laser (HB-Laserkomponenten GmbH, Schwäbisch Gmünd, Germany) was used as the excitation source, and delivered through a 0.65 FCP multiwavelength, polarization-maintaining fibre from Point Source, U.K. The output beam was passed through a dichroic filter (LLF532/10x Brightline, Semrock, Rochester, NY). The laser power was set to ∼1.1 kW cm^−2^. A quarter-wave plate (375/550 nm, B. Halle Nachfolger GmbH, Berlin, Germany) was inserted into the illumination path to ensure nearly circular polarization of the laser beam. The fluorescence emitted by the sample was collected by the same objective lens and separated from the exciting laser light by a dichroic filter (Z580dcxr, AHF Analysentechnik, Tübingen, Germany). Residual excitation laser light was removed by a notch filter (NF01-532U-25, AHF Analysentechnik, Tübingen, Germany) and the detection range was limited to the Texas Red spectrum by a bandpass filter cube (Nikon TRITC).

### Cell-surface fluorescence staining of nAChRs

CHO-K1/A5 cells grown on 18 mm diameter No. 1.5 glass coverslips (WRL) in Ham’s F12 medium at 37°C were washed thrice with M1 medium and incubated for 45 min-1 h at 4°C with mAb35 primary antibody, washed thrice with cold M1 containing 10% bovine foetal serum, and incubated with Texas Red-labelled goat anti-mouse secondary antibody for 1 additional h at 4°C in M1 medium supplemented with 2% bovine serum albumin, washed several times with M1 supplemented with 0.1M glycine and 1% v/v glucose, and mounted in a chamber with STORM buffer (1% v/v glucose oxidase, 1% v/v catalase and 0.1-0,05% v/v 2-mercaptoethanol in M1 medium). Each specimen was imaged live for less than 4 min after exposure to the STORM buffer. In the case of fixed specimens, cells were washed 3x with M1 medium followed by 3x washes with M1 containing 2% BSA, followed by incubation in 4% paraformaldehyde containing 2% sucrose for 10 min at room temperature. After washing several times with M1, fixed specimens were covered with 20 μL of a 1% aqueous solution of 25,000 MW PVA in Millipore-filtered distilled water, spun in a table top centrifuge rotor shaft to form a thin layer of dried PVA (Barrantes 2016), and finally mounted onto round coverslip holder chambers.

### Single-molecule superresolution imaging

Single-molecule localization microscopy in the STORM modality was carried out in a setup constructed in our laboratory. Briefly, a planapochromatic TIRF 100x, 1.49 N.A. oil immersion objective was used in conjunction with a back-illuminated electron multiplying CCD camera (iXon-Plus DU-860, Andor Technology, Belfast, Northern Ireland) set to acquire a stream of images at maximum frame rate at a gain of ∼250, 3.6 photoelectrons per A/D count and pixel size of 106nm. Streaming movies were acquired from between 8 and 15 cells using the software SlideBook (Intelligent Imaging Innovations, Boulder, CO) and exported as 16-bit TIF or Matlab files for subsequent off-line analysis. All images were recorded from the ventral, coverslip-contacting membrane of the cells. Cells were inspected after image acquisition to ensure preservation of cell morphology.

### Superresolution image data analysis

#### i) Determination of sub-diffraction molecular coordinates

The off-line localization of the x,y coordinates of the nAChR spots was carried out using the image analysis package ThunderSTORM (https://code.google.com/p/thunder-storm/) (Ovesný et al. 2014) run as a plugin in ImageJ (https://imagej.nih.gov/ij/). ThunderSTORM is particularly suitable for separating multiple overlapping PSFs (typical emitter density was 5 localizations per frame). In order to account for the discrete nature of pixels in digital cameras, an integrated form of a symmetric 2D Gaussian function was fitted to the spots using Levenberg–Marquardt least-squares minimization routines. Localizations that were too close together to be independent were discarded. The ThunderSTORM multi-emitter fitting analysis was enabled and the limiting intensity range was set at 500-2,000 photons. Other ThunderSTORM filters were enabled to remove uncertainty-based duplicates (e.g. multiple emitters and duplicate localizations, as described in ref. (Huang et al. 2011)). Lateral drift was estimated experimentally using fiducial 100 nm coverslip-adhered fluorescent beads and corrected via the appropriate filter in ThunderSTORM. Localization precision was calculated automatically using a modified version of the formula in ref. (Thompson et al. 2002) which considers the EM gain of the EM-CCD camera (Quan et al. 2010), through the following expression:

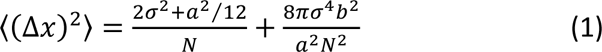

where *σ* is the standard deviation of the fitted point spread function (PSF), *a* is the pixel size in nm, *N* is the intensity expressed in number of photons and *b* is the background signal level in photons. The average localization precision was 40 nm (see Suppl. Fig. 1a).

#### ii) Single-particle tracking (SPT)

The fluorescent particles were detected by a generalized likelihood ratio test algorithm specifically designed to detect point spread function-shaped (i.e. Gaussian-like) spots using ThunderSTORM (Ovesný et al. 2014) and exported in a format suitable for tracking analysis using an ad-hoc Matlab routine written in our laboratory. Detected (validated) particles were further analyzed for their trajectories with the software package Localizer (https://bitbucket.org/pdedecker/localizer), (Dedecker et al. 2012) implemented in Igor Pro (Wavemetrics, Inc. https://www.wavemetrics.com). Two critical parameters were set in Localizer: the maximal number of frames (2 frames = 20 ms) that a given molecule was allowed to blink (“Max blinking”), and the maximum distance (“Max jump distance”) at which two points could lie and be attributed to the same trajectory (2 pixels). The optimal “Max blinking” (t_off_) was determined following the method of Annibale and coworkers (Annibale et al. 2011), which implies that the number of photoblinking fluorescent molecules N in the sample can be estimated from the number of counts (localizations) at different dark times t_d_, N(t_d_), by fitting to the semi-empirical equation:

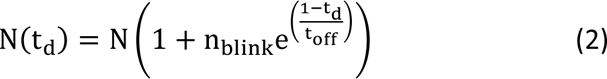

in the regime of low dark time t_d_ values (Annibale et al. 2011). The maximum distance for the merging filter in our experiments was determined from the camera pixel size and the distance-filter criterion (Lu et al. 2014). Briefly, localized molecules that reappeared in consecutive frames were considered as corresponding to the same molecule if the frame-to-frame displacement (tracking radius) was within 106 nm, thus allowing the monitoring of molecules with diffusion coefficients of up to 1.33 µm^2^ s^−1^, i.e. conservatively higher than the upper limit for nAChR nanocluster diffusion estimated from TIRF-SPT experiments in our laboratory (Almarza et al. 2014). Following the above argument, the “Max jump distance” was deduced by considering the maximum distance that a nAChR is allowed to “jump” within the merge time window established above, i.e. the maximum distance that, on average, a single-molecule can travel in 2 frames.

#### iii) Mean-square displacement (MSD) analysis

Typically, a diffusion process is characterized by the mean-square displacement (MSD), which for a 2-dimensional space like a membrane bilayer can be written as:

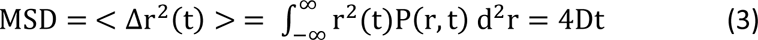

where D is the diffusion constant. This assumes a purely viscous and homogeneous fluid, such that P(r, t) is the probability distribution function (PDF, also termed propagator) of the diffusion process, i.e., the probability of finding the particle at a (radial) distance r away from the origin at time t after release of the particle at r = 0 at time t = 0.

Complex media may lead to sublinearity of the MSD as a function of time:

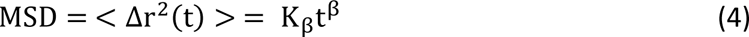

In equation 4, anomalous diffusion is taken into account by introduction of the exponent β (Tejedor et al. 2010; Manzo et al. 2015), where (MSD) ∼ t^β^. Whereas for β = 1 simple thermally-driven (“random walk”) Brownian diffusion results, two forms of anomalous diffusion result from other values of β: subdiffusion for 0 < β < 1 (e.g. in molecular crowding), and superdiffusion for β > 1 (i.e. motor-driven transport). Thus, in the case of anomalous diffusion Eq. (3) above can be written in short-form as: 4Dt^β^.

The time-averaged mean-square displacement (TA MSD) was calculated for each trajectory j in the form:

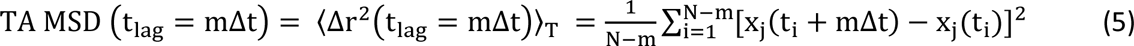

where x_j_ is the position sampled at N discrete times t_i_ = iΔt, Δt the acquisition time (in our case, Δt = 10 ms) and i the frame number.

The ensemble-averaged mean-square displacement (EA MSD) was calculated over a time interval mΔt,

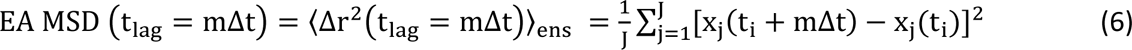

where J is the total number of single-molecule trajectories (not NaN) at time t_i_, and t_i_ is the starting time relative to the first point in the trajectory.

The ensemble average of the temporal-averaged mean-square displacement (EA TA MSD) was calculated by ensemble averaging the TA MSD or by temporal averaging the EA MSD, by truncating the data at different observation times T:

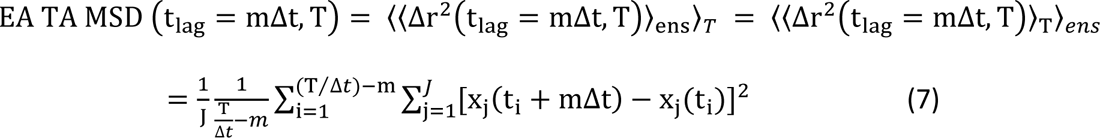

Here we followed the nomenclature employed by Krapf and coworkers and TA, EA and EA TA MSD were calculated in accordance with their procedures (Weigel et al. 2011). The 95% confidence interval for the EA MSD was calculated following the criteria of ref. (Janczura and Weron 2015). Following the criteria of ref. (Golan and Sherman 2017) the anomalous exponents (β) were obtained by linearly fitting the initial 50 points of the log-log transformed TA MSDs. The generalized diffusion coefficients (*K_β_*) were obtained from the linear fit to the first 50 time points of the individual MSDs in log-log scale, evaluated at *t* = 1 (see Eq. 4 above), also following the criteria of (Golan and Sherman 2017).

#### iv) Exclusion of immobile particles from the analysis of nAChR trajectories

Stationary (immobile) molecules were excluded from the analysis of single-molecule trajectories following a series of recently introduced criteria (Golan and Sherman 2017). The procedure sets a threshold value on the ratio of the radius of gyration R_g_ and the mean step size |Δr| of the particles’ displacement. In the case of ideal immobile particles this ratio is constant, whereas for mobile particles the ratio increases. The normalized ratio:

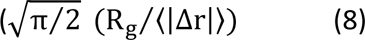

was obtained from experiments with 4% paraformaldehyde-fixed cells, and the ratio was subsequently employed to obtain the threshold value applied to live cell experiments. Golan and Sherman (2017) discuss the advantages of this method over the use of the diffusion coefficient or R_g_ alone for excluding immobile particles; the two latter procedures would falsely classify immobile particles as mobile. Threshold values with 95% confidence were obtained by pooling data from different cells in independent sets of experiments.

#### v) Turning angle analysis

The turning angle distribution as a function of the time lags has been employed to search for correlations in the particleś displacements (Burov et al. 2013) (Sadegh et al. 2017). We applied this analysis to follow the directional changes in individual nAChR trajectories using the relative angles distended by the molecules along their walk in successive time intervals, a parameter which can help distinguish among different types of anomalous subdiffusive mechanisms, e.g. the so-called fractional Brownian motion (fBM) and obstructed diffusion (OD) models.

#### vi) Single-molecule trajectory recurrence analysis

The presence of confinement periods within individual trajectories was identified using recently developed algorithms (Sikora et al. 2017), based on the evaluation of the total number of visits (recurrence) performed by the moving particle to a given site. Confinement within a nanoscale domain is associated with visits to the same site, multiple times within a short period, whereas unconfined motion is related to exploration of wider regions, less compact walks, and less visits to the sites previously walked.

#### vii) Identification of dynamic nAChR nanocluster formation/disassembly

We applied the centroid-linkage hierarchical clustering and the density-based spatial clustering with noise (DBSCAN) analyses embedded in the open source qSR software developed by Cissé and coworkers (Andrews et al. 2018) (www.github.com/cisselab/qSR) to identify in real time the formation and disassembly of clusters of nAChR molecules. The length scale was set at 100 nm, and the minimal number of particles was 10 in the DBSCAN analysis. Dark-time tolerance was set at 1 s. For this purpose and only in this analysis, the immobile nAChRs were included.

### Statistical analyses

These were done using one-way analysis of variance employing Kruskal-Wallis or Sidak’s multiple comparisons test in the GraphPad Prism 6 software. Mean ± 95% confidence interval is shown unless otherwise stated. The one-sample Kolmogorov-Smirnov test was applied to assess whether the data were normally distributed or not. To compare two distributions, we used the Kolmogorov-Smirnov (KS) test for two samples. The experimental procedures (cell cultures, immunocytochemistry, STORM imaging) and the statistical analyses were performed by different individuals.

## Results

### Antibody tagging results in a high degree of nAChR immobilization

CHO-K1/A5 cells, a clonal cell line robustly expressing adult muscle-type nAChR (Roccamo et al. 1999) were used throughout. Live CHO-K1/A5 cells labeled with the monoclonal antibody mAb35 against the nAChR α-subunit followed by Texas Red-labeled secondary antibodies were imaged at their coverslip-adhered (ventral) surface under control or cholesterol-modifying conditions at a frame rate of 100 frames per second as detailed in Material and Methods. Between 8 and 15 cells were recorded per condition. Reconstruction of the superresolution images and single-molecule tracking were done using the ThunderSTORM software (Ovesný et al. 2014) and the Localizer package (Dedecker et al. 2012)). In order to ensure the robustness of the results, trajectories shorter than 50 temporal steps (i.e. with a difference of 50 frames between the initial and the final recorded positions) were discarded. However, an important fraction of the remaining trajectories displayed a high level of confinement. The possibility that these trajectories correspond to immobile receptors exhibiting an apparent movement due to the localization error cannot be totally discarded. Since this could affect the results and lead to erroneous conclusions about the diffusional properties of the ensemble, we implemented the criteria described by Golan and Sherman (Golan and Sherman 2017), based on the ratio of the radius of gyration and the mean step size. An important advantage of this procedure is its independence from the localization error. The ratio is nearly constant for immobile receptors displaying an apparent motion but increases for mobile receptors. We calculated the distribution of this ratio for apparent trajectories of completely immobile receptors from fixed specimens, and the upper 95% confidence interval was chosen as the cutoff for the nAChR molecules in live cells. Trajectories with lower ratios were considered immobile and not included in the diffusional analyses. A total of 80 ± 9 % immobile nAChRs was obtained under control conditions, as already apparent by visual inspection of the individual tracks, with no significant changes upon cholesterol modification (Suppl. Fig. 2).

### Antibody-crosslinked nAChRs display ergodic-like heterogeneous diffusion

Mean squared displacement (MSD) is the most widely used criterion to detect deviations from random (Brownian) motion in protein diffusion analysis. Typically, it is studied by a temporal-average over time (TA MSD) for single-particle trajectories or by an ensemble-average over several particles for the population of trajectories at a specific time (EA MSD). The nonlinear fit of the individual TA MSD yields both the anomalous exponent (β, a measure of the deviation from Brownian motion) and the generalized diffusion coefficient (K_β_) (see relevant equations in Material and Methods. A β value different from 1 is associated with anomalous (non-Brownian) diffusion: β<1 for sub-diffusion (attributed to protein-protein immobilizing interactions, confinement, or spatial inhomogeneities) and β>1 for super-diffusion (usually resulting from active cellular transport processes). At the single-molecule level, nAChR dynamics was heterogeneous: from subdiffusive to Brownian and seldom superdiffusive. The averaged β values of the population were below unity, i.e. subdiffusive, and averaged generalized diffusion coefficients K_β_ were in the order of 0.5 μm^2^s^−β^ (Figure 1a). No significant changes were observed in these two parameters upon modification of cholesterol membrane content (Suppl. Fig. 3).

**Figure 1.**
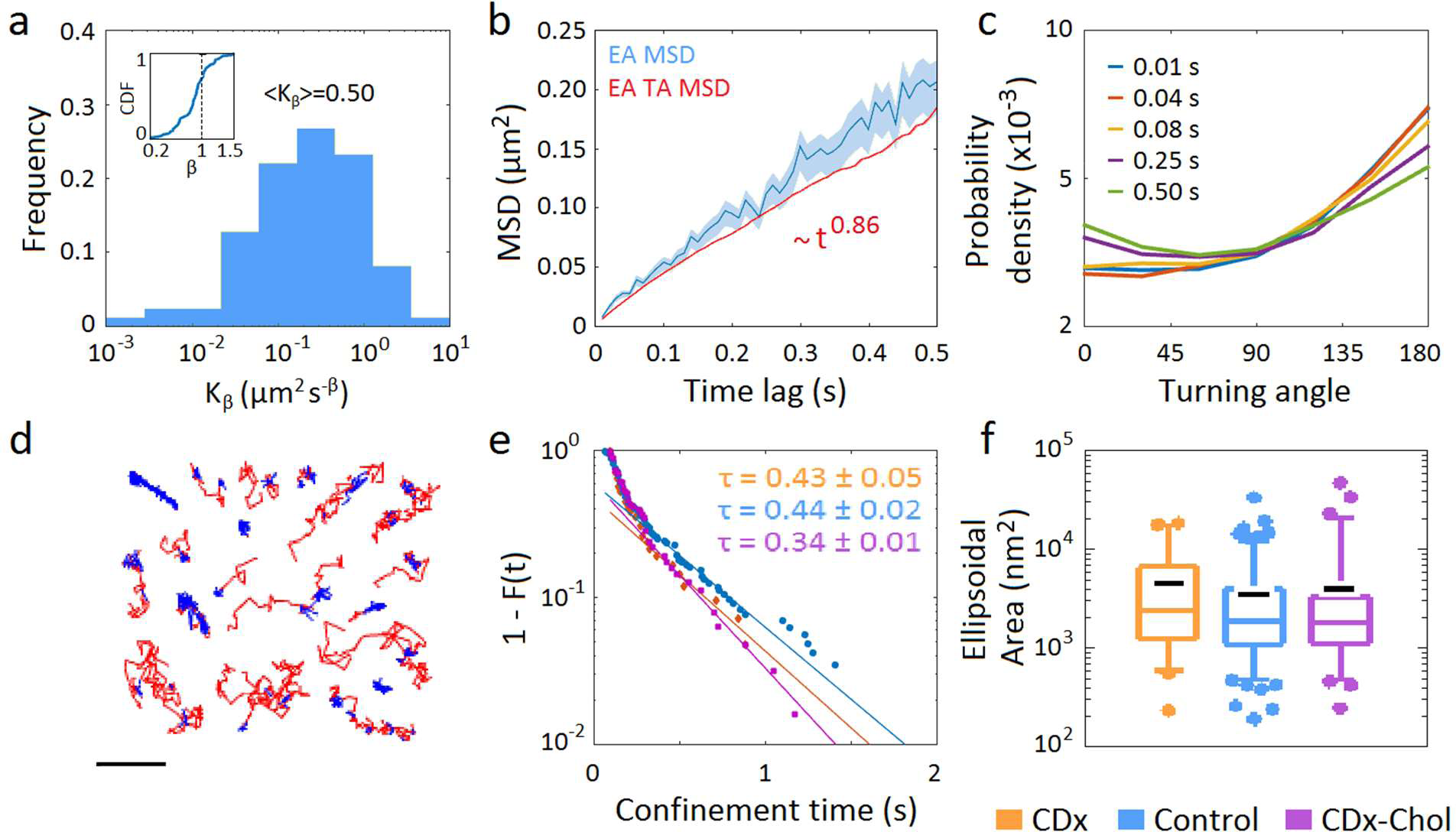
Mean-square displacement, turning angles and recurrence analysis applied to single-molecule nAChRs under control and cholesterol-modifying conditions. a) Frequency of the generalized diffusion coefficients (K_β_) in log scale and cumulative density function of the anomalous exponent β (inset), obtained by linear regression of the temporal-averaged MSD (TA MSD) in log-log scale of single-molecule trajectories (see Methods for details) (n = 149 trajectories). No changes were observed after modifying the membrane cholesterol content (Suppl. Fig. 3). b) The ensemble-averaged MSD (EA MSD) with the 95% confidence interval is shown in blue. An additional ensemble averaging to the TA MSD (EA TA MSD, red trace) was performed to test the ergodic hypothesis. Similar trends were observed for cholesterol-modifying conditions (see Suppl. Fig. 4). c) Probability densities of the turning angles for the (color-coded) time lags (t_lag_) of increasing durations (10 to 500 ms). The probability density is normalized such that the integral of the curve is equal to unity. For cholesterol-modifying conditions see Suppl. Fig. 5. d) Examples of the results obtained by applying the recurrence analysis (Sikora et al. 2017) to the single-molecule nAChR trajectories. The confinement sojourns are shown in blue. Scale bar: 1 μm. e) Complementary cumulative distribution function of the resident times in the confined state for the nAChRs under control (blue circles), cholesterol-depletion (orange diamonds) and cholesterol-enrichment (purple squares) conditions. The distributions were exponentially tailed, with the characteristic decay times (τ) stated in the figure (in seconds). f) Ellipsoidal area of the trajectorieś portion in the confined state, obtained by fitting the confinement domain in the trajectory with an elliptic surface. Whiskers in box plots correspond to 95% confidence intervals; the limits indicate 75% confidence intervals; the black minus symbols indicate the mean and the colored horizontal lines the median in each case. The dots outside the confidence intervals are outliers.

Single-particle analyses often require calculating temporal averages over individual trajectories from which the diffusional properties of the ensemble are obtained. Yet this is correct only under the assumption that the individual molecules can explore all accessible diffusion states within the recording time, fulfilling the ergodic hypothesis. If this assumption does not hold, the diffusional properties obtained by temporal averaging over a single molecule will not be equivalent to those resulting from ensemble averaging over the entire population. Many biological systems conform to this latter behavior and hence display weak ergodicity breaking (WEB), often related to transient immobilization or spatial inhomogeneities (Weigel et al. 2011; Manzo et al. 2015; Weron et al. 2017). For this reason, testing the ergodic hypothesis can unravel important features of the diffusion of the protein in the membrane. Classically, the ergodic hypothesis is assessed by comparing the TA MSD and the EA MSD. In theory, if the system is ergodic, the TA MSD converges to the EA MSD and the parameters estimated from both averages (e.g. anomalous exponent) will be equal. However, the empirical and the theoretical values will always differ (especially for small samples), making it necessary to consider a measure of the accuracy of the empirical estimation (e.g. including the confidence interval) without which a process could be falsely classified as non-ergodic due to sample size limitations. Following this idea, we compared the EA TA MSD (ensemble average of the TA MSD) and the EA MSD together with the 95% confidence interval, as implemented in ref. (Janczura and Weron 2015). In our system, the curves of the EA TA MSD and the EA MSD appear to reflect an ergodic-like behavior (Figure 1b and Suppl. Fig. 4).

### nAChRs exhibit anticorrelated steps that conform to diffusion in a medium with a characteristic size and high obstacle density

We next attempted to elucidate the mechanism behind the anomalous but ergodic diffusion of the nAChR. A simple way to do this is by inspecting the distribution of the directional changes or turning angles followed by the molecules, i.e. the relative angles distended by successive molecular “walks” for different time intervals (Burov et al. 2013). If the particles do not show any preferred direction of motion (i.e. in Brownian motion), the turning angle distribution is expected to be flat; instead, the distribution peaks at 0° if they exhibit a preference to move forward or at 180° when the preferred motion is backwards. The turning angle distributions of the nAChR at the cell surface increased from 60° to reach their maximum at 180° (Figure 1c), a clear manifestation of subdiffusive random walks with anticorrelated steps (i.e. with a predilection of individual molecules to retrace their steps). By numerical simulations, Krapf and coworkers analyzed the turning angle distributions of two widely tested ergodic biophysical models (Sadegh et al. 2017) that led to subdiffusion with anticorrelated steps: motion of particles in a viscoelastic fluid (fractional Brownian motion) and motion hindered by nearly immobile obstacles (obstructed diffusion). Although the turning angle distribution peaked at 180° in both models, in the first one it reached a plateau, in clear contrast with the present experimental data, indicating that nAChR dynamics is better explained by the obstructed diffusion model. Moreover, two additional properties can be obtained from inspection of the turning angle distributions. Firstly, by comparing the abruptness of the present distributions with those obtained for simulations of particles undergoing obstructed diffusion with increasing obstacle concentrations (Sadegh et al. 2017) we can infer that nAChR diffusion is hindered by a relatively high concentration of immobile particles. Secondly, we observe a significant dependence of turning angles with time lag (statistical differences between t_lag_ = 0.01 s and t_lag_ = 0.50 s were *p* < 0.01, for cholesterol depletion, *p* < 10^−1^, for control and *p* < 10^−5^, for cholesterol-enrichment conditions, respectively) (see Suppl. Fig. 5). This feature is related to anticorrelated diffusion in a structure with a characteristic size, as was previously observed in numerical simulations of random walks in a permeable fence model (Sadegh et al. 2017), an observation which enables us to discard the involvement of fractal structures. That is, nAChR molecules move by random walks within a meshwork with a low probability of permeation. In summary, our turning angle data conform to the general depiction of molecules diffusing in a structure with a characteristic size constituted by immobile obstacles at a high concentration.

### Transient confinement nanodomains are a characteristic feature of nAChR motion

If receptors diffuse in a medium with a characteristic size and high obstacle density-which implies a low probability of permeation-one would expect to observe motional hindrance in this medium. When a particle is confined, it tends to visit the same area repeatedly until it eventually escapes. Several mechanisms have been proposed to account for these trapping events (e.g. molecular crowding, protein-protein interactions, interactions with the actin cytoskeleton and membrane heterogeneities). A recently proposed analysis (Sikora et al. 2017) allows one to identify regions of confinement in individual trajectories, and extract the characteristic sojourn durations in the recurrently-visited regions where the particle remains trapped. When applied to the nAChR trajectories, we observed an interesting phenomenon: essentially all trajectories (from subdiffusive to Brownian) alternate between free walks and confinement sojourns with a well-defined length scale (see Figure 1d). The confinement zones could therefore potentially be the structure described by the turning angles analysis. The residence time distribution in the confined state was exponentially tailed, with characteristic decay times of 430 ± 50 ms, 440 ± 20 ms and 340 ± 10 ms, for cholesterol depletion, control, and cholesterol enrichment conditions, respectively (Figure 1e). The average surface distended by a confinement sojourn had a radius of ∼34 nm (Figure 1f), a value which was not significantly affected by changes in membrane cholesterol levels.

When we repeated the turning angle analysis on the confined and free portions of the single-molecule trajectories separately, we realized that the anticorrelated steps occurred only within the confined areas, whereas the free walk periods displayed a nearly flat Brownian distribution (Figure 2). *This confirms the idea that the confinement zones are the structures behind the anticorrelated steps.* An individual receptor molecule would thus appear to move freely across the cell membrane until it encounters a region of high obstacle density that hinders its movement for a certain period until it escapes.

**Figure 2.**
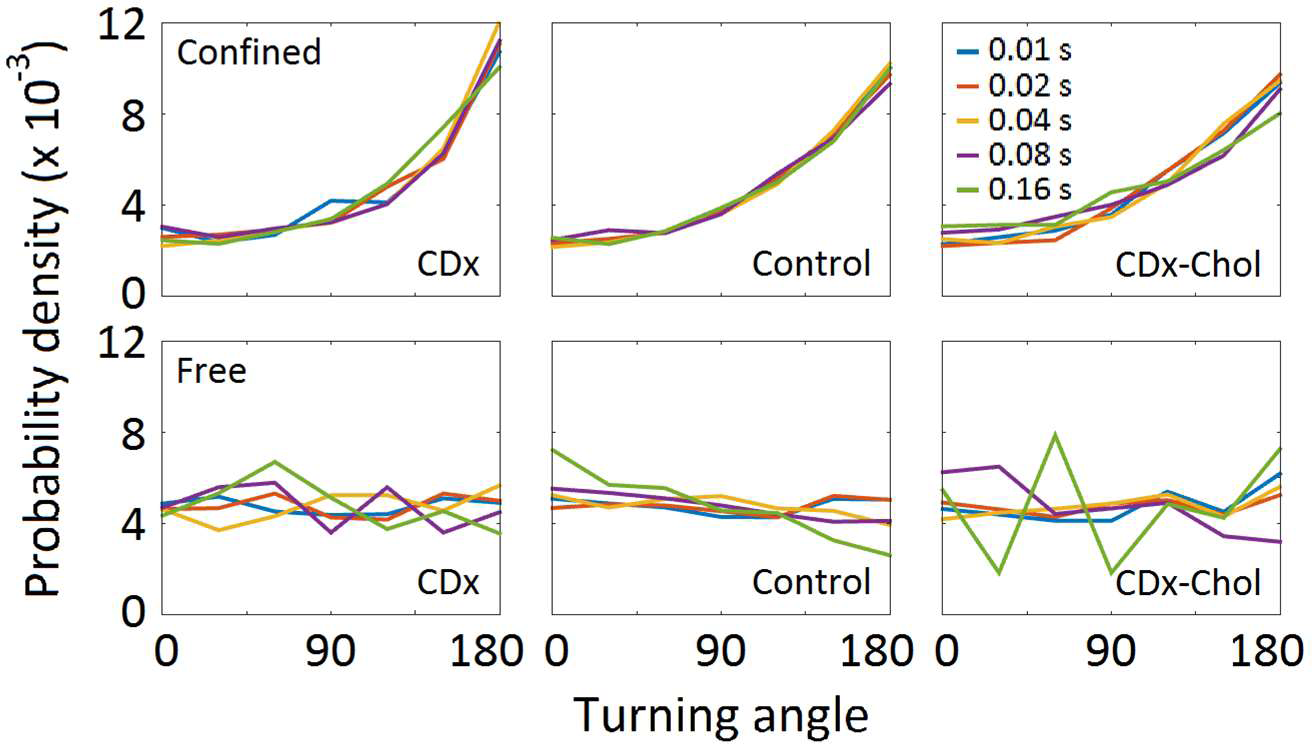
Turning angle probability densities of the transiently confined (upper row) and unconfined (free; bottom row) portions of the individual trajectories under control and cholesterol-modifying conditions. Probability densities for the color-coded t_lag_ of increasing durations (10 to 160 ms) (n = 114-392 confined trajectories and n = 124-412 free trajectories). The probability density is normalized such that the integral of the curve is equal to unity.

### Clustering event dynamics and extension are modulated by membrane cholesterol levels

To address the possible physical nature of the immobile obstacles that hinder receptor mobility in the confinement sojourns, we examined the dynamics of cluster formation/disassembly. Using the qSR software developed by Cisse and coworkers (Cisse et al. 2013; Andrews et al. 2018), we were able to follow the temporal evolution of clustering events in live CHO-K1/A5 cells. Briefly, this analysis allows one to identify spatial clusters from superresolution data and extract key descriptors such as duration, area and number of localizations per cluster. nAChR trajectories display clustering events (assembly) with an average duration of 11 ± 1 s (Figure 3a), typically followed by cluster disassembly periods. The intra-burst mean dark time (i.e. the time between two particle localizations within a clustering event) was significantly affected by cholesterol depletion, increasing from an average of 241 ± 6 ms to 274 ± 10 ms (Figure 3b). The radius of the zones where the clustering events occur was inversely proportional to cholesterol concentration, the control radius of 139 ± 6 nm increasing to 182 ± 7 nm upon cholesterol depletion and decreasing to 118 ± 7 nm with cholesterol enrichment (Figure 3c). The number of single molecules within an assembly was 50 ± 6, decreasing in a statistically significant manner to 46 ± 9 after cholesterol depletion (Figure 3d).

**Figure 3.**
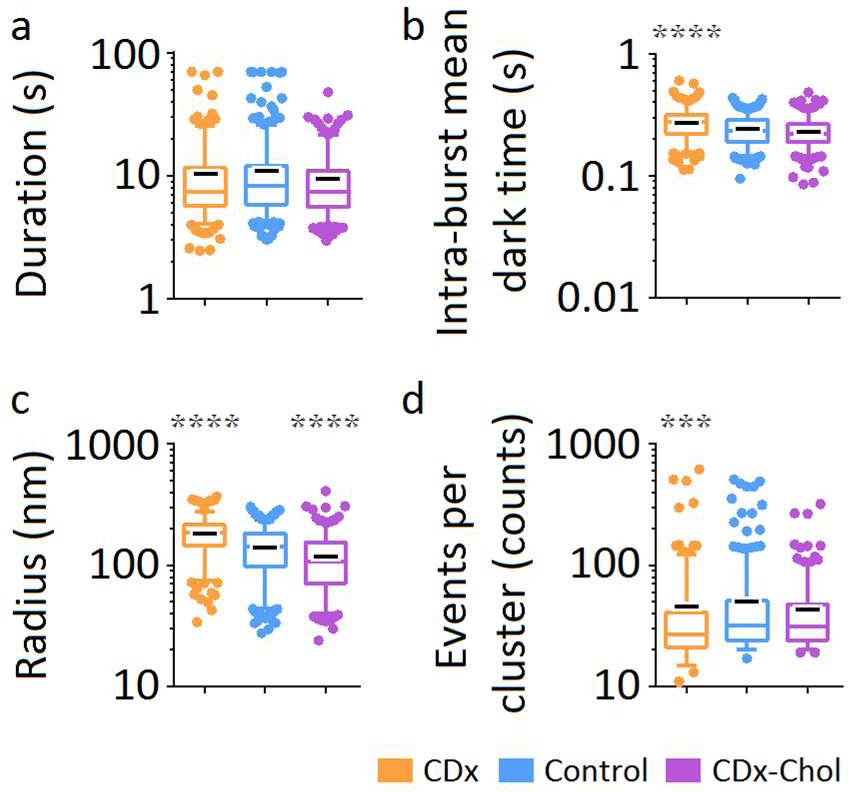
Real time follow up of nAChR dynamic clustering and disassembly under cholesterol depletion and enrichment. Using the live cell single-molecule localizations validated with ThunderSTORM as the input data, the qSR software (Andrews et al. 2018) was applied to obtain the nAChR clustering metrics (log scale): duration (a), intra-burst mean dark time, i.e. periods without localizations within a clustering event (b), radius (c), and number of occurrences per clustering event (d) under control and cholesterol-modifying conditions (n = 206-387 clustering events). Whiskers in box plots correspond to 95% confidence intervals; the limits indicate 75% confidence intervals; the black negative symbols indicate the mean and the colored horizontal lines the median in each case. The dots outside the intervals are outliers. Statistics: *p* < 0.001 (***) and *p* < 10^−4^ (****).

## Discussion

### Antibody-mediated crosslinking dramatically increases receptor immobilization

The tightly-packed nAChRs at the adult NMJ are essentially immobile for < 8 hr (Akaaboune et al. 2002). Labeling with the divalent antibody mAb35 crosslinks nAChRs, resulting in a very high proportion of immobile molecules (∼80%) (Suppl. Fig. 2). In contrast, using the fluorescent monovalent ligand Alexa Fluor^532^ α-bungarotoxin (BTX), immobile receptors represented ∼50% of the total population (Mosqueira et al. 2018). The possibility that this lower figure is caused by the unbinding of BTX from the receptor is unlikely, given that the dissociation of fluorescently-tagged BTX from the nAChR at a live NMJ takes between 9 and 14 days. In other words, the disappearance of BTX-bound receptors is mainly due to nAChR turnover and not unbinding (Akaaboune et al. 1999).

The translational diffusion of the remaining mobile mAb-tagged nAChR ensemble population is heterogeneous and mostly subdiffusive. Remarkably, the generalized diffusion coefficient of the ensemble is higher in mAb-labeled nAChRs than in BTX-labeled nAChRs in both control and cholesterol-depleted conditions (*p* < 0.05) (this work and ref. (Mosqueira et al. 2018)). The above-mentioned work of Akaaboune and coworkers identified a “dynamic fraction” of receptors at the mature NMJ. Considering the dramatic increase of receptor trapping and immobilization, the augmented lifetime of the mobile receptors in the confined state and the increased duration of the clustering events observed here in a model mammalian cell, it is possible that the mobile fraction of receptors in the mature NMJ described by Akaaboune and coworkers constitutes a preferential target of antibody binding and crosslinking in myasthenia and related pathologies.

### mAb35-crosslinked nAChRs have longer confinement sojourns than BTX monovalently-tagged receptors

Turning angle analysis revealed anticorrelated steps, indicating that nAChRs are more likely to retrace their steps than to move forward (Figure 1 and Suppl. Fig. 5). Time lag dependence was also observed, in good agreement with diffusion in a structure with a characteristic length scale, as was observed by numerical simulations of particles diffusing in a permeable fence model (Sadegh et al. 2017). Moreover, dissecting all trajectories into confined and free (unconfined) portions allowed us to disclose some important features of the nAChR translational behavior: it is the confined portion of the trajectories that exhibits turning angle distributions with anticorrelated steps, thus conforming with the obstructed diffusion model, whereas the nearly flat distributions of the free portion of the walks correspond to Brownian diffusion (Figure 2). Thus, moving mAb-labeled nAChRs display free Brownian displacement across the cell membrane, intermittently interrupted by confinement sojourns with a well-defined length-scale, within regions of high obstacle density, that is conforming to a two-state switching diffusion model (Grebenkov 2019) with residence times comparable to the length of the trajectories. Antibody-tagged receptors exhibit longer (340-440 ms) confinement sojourns than those labeled with a monovalent ligand like α-BTX (135-257 ms) (Mosqueira et al. 2018), indicating that mAb-crosslinked receptors remain for longer periods in confined areas.

### Antibody cross-linking stabilizes/prolongs the cholesterol-dependent dynamic nanoclustering events

In addition to the brief stopovers (transient confinement sojourns) of individual trajectories (Figure 1d), nAChR molecules also undergo a collective dynamic clustering phenomenon whereby they assemble and disassemble in a strongly cholesterol-sensitive manner, in areas roughly 4-7-fold larger than the individual confinement areas (Figure 3c). Indeed, cholesterol depletion increases the size of these larger assembly/disassembly zones (from radii of ∼140 nm under control conditions to ∼180 nm), whereas cholesterol enrichment reduces radii to ∼120 nm (Figure 3). When the mAb-labeled nAChR clustering events are compared with those of BTX-labeled receptors (Mosqueira et al. 2018) the lifetimes of the antibody nanoclustering events are found to be 3-fold longer (p < 0.01 for control and *p* < 10^−4^ for cholesterol-modifying conditions), a finding consistent with the idea that mAb35-induced crosslinking adds stability to/prolongs the nanocluster formation. These dynamic structures may correspond to the diffraction-limited “microaggregates” observed along NMJ development (Olek et al. 1986a; Olek et al. 1986b) reviewed in (Sanes and Lichtman 2001).

### Implications of mAb35-nAChR complex diffusion in the framework of myasthenia

Because of their crosslinking ability, mAbs against the MIR of the nAChR share the capacity to activate the physiopathological events induced by myasthenic autoantibodies (see recent reviews in (Gilhus 2016; Paz and Barrantes 2019). These processes include mainly a complement-mediated focal lysis of the postsynaptic membrane (Drachman et al. 1978; Engel and Arahata 1987) that together with the so-called antigenic modulation (Conti-Tronconi et al. 1981)which enhances endocytic internalization of the receptor, produces a significant reduction in cell-surface nAChR numbers in myasthenia. We have not only shown that the endocytic process is effectively triggered by divalent mAb35 molecules in CHO-K1/A5 cells (Kumari et al. 2008) but also that it is significantly accelerated upon cholesterol depletion (Borroni and Barrantes 2011). As shown in the present work, mAb35-mediated crosslinking results in a very high proportion of immobile receptors, while the remaining mobile ones undergo anomalous ergodic diffusion governed by stopovers within zones of high obstacle density. Cholesterol modulates receptor diffusion, and its depletion from the membrane increases the size of the dynamic receptor clustering events. The latter may further increase nAChR degradation.

About 80% of patients with generalized myasthenia gravis have serum antibodies to the nAChR (Vincent 2002). Patients with neither anti-nAChR nor anti-MusK autoantibodies are called seronegative, and one of the hypotheses on the apparent seronegativity is that the autoantibodies of these patients react only with the membrane-associated and clustered form of the nAChR (Leite et al. 2008) as found in the native neuromuscular junction at very high densities. Here, we show that antibody-mediated nano-clustering already suffices to affect nAChR mobility.

What is the possible relationship between the observed cholesterol modulation of nAChR diffusion and myasthenia? Although the evidence is scanty, it has been reported that 3-hydroxy-3-methyl glutaryl coenzyme A reductase inhibitors, commonly known as statins and used therapeutically by inhibiting cholesterol biosynthesis, may trigger anti-nAChR autoantibody production (Purvin et al. 2006), and exacerbate or unmask subclinical myasthenia gravis, most often reversibly (Cartwright et al. 2004; Purvin et al. 2006) (Oh et al. 2008), with increases in anti-nAChR autoantibody titers (reviewed by (Gilhus 2009). We also know that cholesterol depletion reduces the number of nAChRs by accelerating the rate of endocytosis and shifting the internalization to a different endocytic pathway involving the small GTPase Arf6 (Borroni and Barrantes 2011). Moreover, we observe in this work that cholesterol depletion increases the size of nAChR clustering events. Thus, these two processes, antibody-induced clustering and cholesterol depletion-enhanced endocytosis, appear to act synergistically to ultimately decrease the cell-surface nAChR population.

Summarizing, at the individual molecule level the nAChR undergoes complex motional regimes caused by a two-state switching between free (Brownian) walks and confined diffusion within zones of high obstacle density acting as permeable fences (∼34 nm radius) (Figure 4). At the ensemble level nAChRs undergo dynamic clustering, trapping and immobilization in much larger areas (120-180 nm radius), which strongly depend on membrane cholesterol levels, suggesting that cholesterol-rich platforms are involved in nanocluster assembly and disassembly. The larger size of the antibody-crosslinked receptor nanometer-sized assemblies may additionally contribute to the immobilization of the protein. This is a clear example of how unraveling the biophysical properties of a key neurotransmitter receptor can throw light on the physiopathological basis of a disease condition, in this case the antigenic modulation characteristically observed in the NMJ affected by autoimmune disease myasthenia gravis.

**Figure 4.**
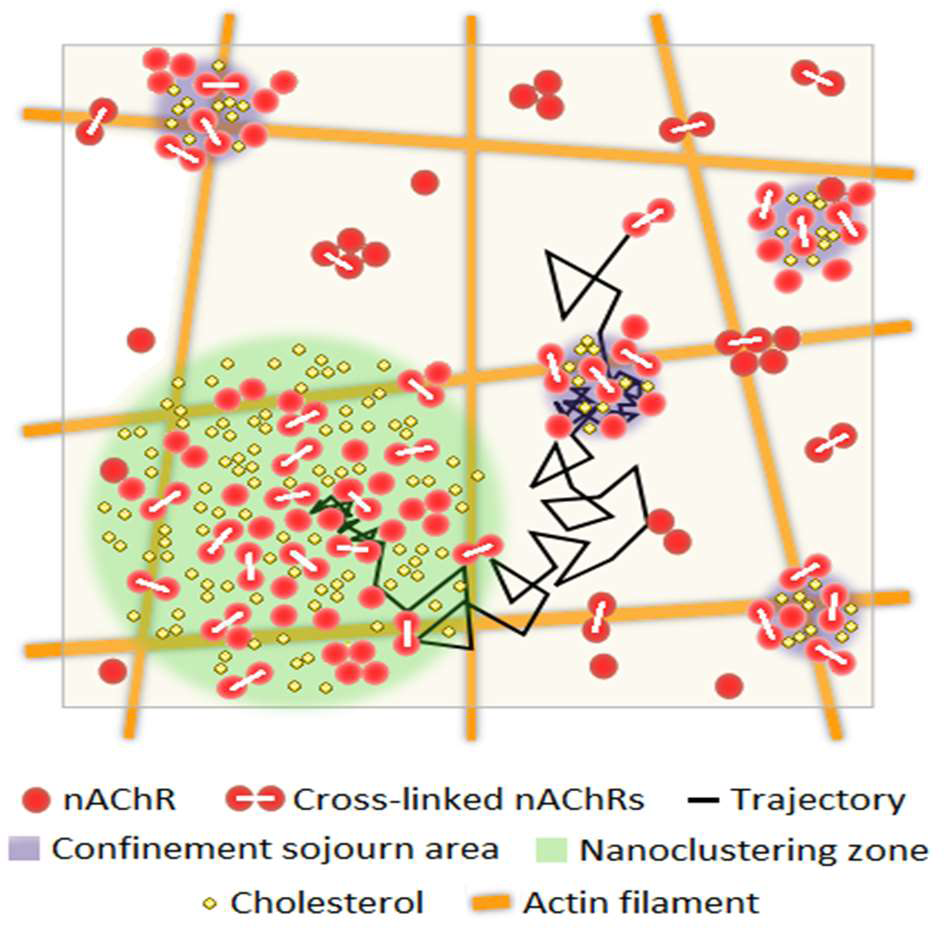
Schematic diagram depicting the complex translational motion followed by nAChR molecules at the cell surface. At the individual level, a nAChR macromolecule (red dots) exhibits a two-state switching behavior between free (Brownian) walks interrupted by transient confinement sojourns. These interruptions occur within small (∼34 nm radius) areas of high obstacle density (purple circles) where receptors transiently reside for an average of ∼400 ms. Dynamic clustering, trapping and transient immobilization also occur in larger nanoclustering zones (∼150 nm radius, green circle) with longer lifetimes (11 ± 1 s), in a strongly cholesterol-sensitive manner. Cholesterol depletion increases the size and average duration of the nanoclustering phenomenon; cholesterol enrichment has the opposite effect. The cortical, sub-membrane actin meshwork may also act as a corrals or fences hindering nAChR diffusion.

## Acknowledgments

We thank Dr. Diego Krapf (Univ. of Colorado) for critical reading of the ms. Experimental work was supported by grant PIP 5205/15 from the National Scientific and Technical Research Council of Argentina (CONICET) to F.J.B.

