## Supplementary material for "Antibody-induced crosslinking and cholesterol-sensitive, anomalous diffusion of nicotinic acetylcholine receptors": Suppl. Material

### Localization precision, merge time and emitter density

The localization precision, merge time and emitter density are three essential parameters in STORM and in single-molecule localization microscopy in general. Suppl. Figure 1 shows the histograms from which the localization precision and the typical emitter density of mAb-labeled nAChR molecules were obtained, as well as a representative example of the merge time distribution and the semi-empirical prediction.

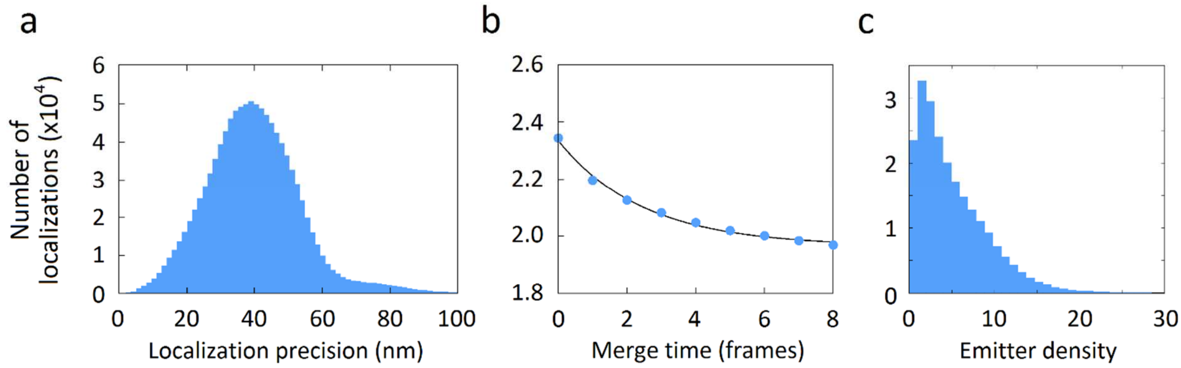


**Supplementary Figure 1. Localization precision, merge time and emitter density**. a) Histogram of the localization precision using the method of Nienhaus and coworkers (Quan et al. 2010). The average localization precision was 40 nm. b) Determination of the optimal merge time following the method of Annibale and coworkers (Annibale et al. 2011). The plot shows the total number of localizations against the merge time in a representative image of mAb-labeled nAChRs and the corresponding fit to the semi-empirical equation (2). In the example shown, the optimal merge time was found to be two frames (20 ms). c) Histogram of the number of localized molecules per frame, i.e. the emitter density. The mean emitter density was 4.70 ± 0.01 molecules per frame.

### Classification of nAChR tracks into mobile and immobile trajectories

The ratio of the radius of gyration $R_g$ and the mean step size can be successfully combined to discriminate between mobile and immobile single molecules (Suppl. Figure 2). A cutoff value of 2.04 was used to exclude immobile molecules (threshold set for 5% false positives), which were omitted in subsequent analyses. Given the lack of non-receptor scaffolding proteins in the CHO-

K1/A5 mammalian clonal cell line, immobilization must respond to the self-aggregation of the nAChR protein in higher oligomeric species (Barrantes 1982), to the crosslinking resulting from antibody binding, or a combination thereof.

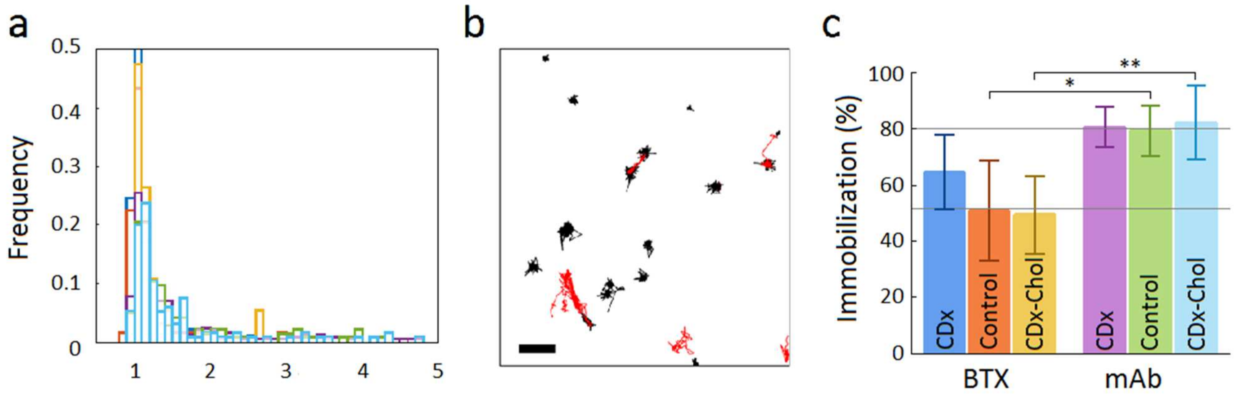


**Supplementary Figure 2. Criteria for discrimination of nAChR trajectories into mobile and immobile.** a) Distribution of the ratio of the radius of gyration and the mean step size for two independent cell culture preparations of nAChRs in paraformaldehyde-fixed cells (6 cells). Threshold values of 1.5-2.04 (95% confidence) were obtained and the conservative value of 2.04 was chosen. b) Mobile (red) and immobile (black) nAChR trajectories under control conditions resulting from application of recently established criteria (Golan and Sherman 2017) for separating particle walks into these two categories. Scale bar: 500 nm. c) Comparison of the percentages of immobile α-bungarotoxin (BTX)- (Mosqueira et al. 2018) and mAb-labeled nAChR receptors (this work) upon application of the threshold resulting from Suppl. Figure 2a; control vs cholesterol depletion with CDx and cholesterol enrichment with CDx-Chol complex. Bars indicate mean ± S.E.M. Statistics: $p < 0.05$ (*) and $p < 0.01$ (**).

**MSD analysis of single-molecule trajectories**

MSD analysis allows one to study the diffusional motifs of single molecules or the ensemble behavior of receptor dynamics at the cell membrane. Linear fits to the log-log transformed temporal-averaged mean square displacement (TA MSD) of individual nAChR trajectories yielded both the anomalous exponent (β) and the generalized diffusion coefficient ($K_\beta$), as defined in eq. 4. Only those linear regressions with a goodness of fit better than 0.9 were included in this analysis. Our results show a heterogenous mobile population, with subdiffusive, Brownian and superdiffusive trajectories. The majority of the nAChRs trajectories were subdiffusive. The average generalized diffusion coefficients were in the order of 0.5 $\mu m^2 s^{-\beta}$. We did not observe statistically significant differences between the experimental conditions for the anomalous exponent or for the generalized diffusion coefficient (see Supplementary Figure 3).

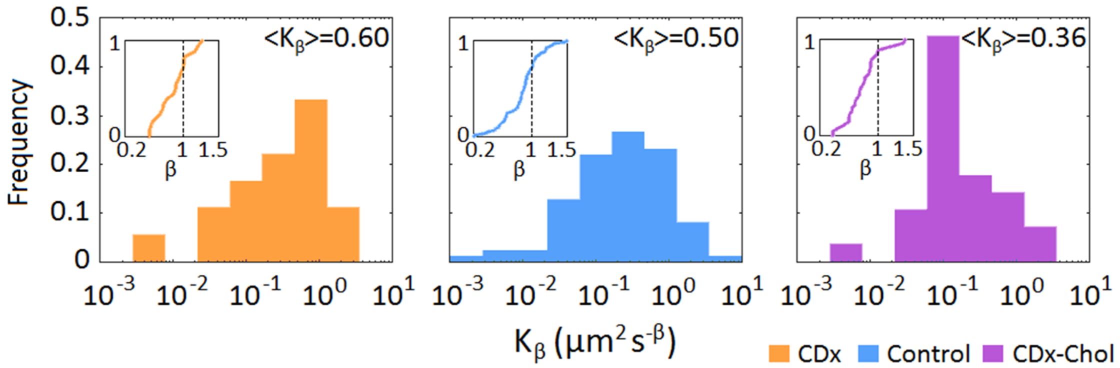


**Supplementary Figure 3.** Normalized distribution of the generalized diffusion coefficients for mAb-labeled nAChRs under control and cholesterol-modifying conditions (8-15 cells, 55 to 149 trajectories). Mean values are shown in each case. The insets show the cumulative density function of the anomalous exponents for each condition. The dotted lines are visual aids corresponding to Brownian motion ($\beta = 1$).

**Ergodicity analysis**

The ergodic hypothesis assumes that the TA- and EA MSD are equivalent for sufficiently long recording times. One way to robustly test this hypothesis consists of comparing the EA TA MSD and the EA MSD together with the 95% confidence interval (see (Janczura and Weron 2015)). The antibody-bound nAChR displays an ergodic-like behavior under control and cholesterol-modifying conditions (see Suppl. Figure 4).

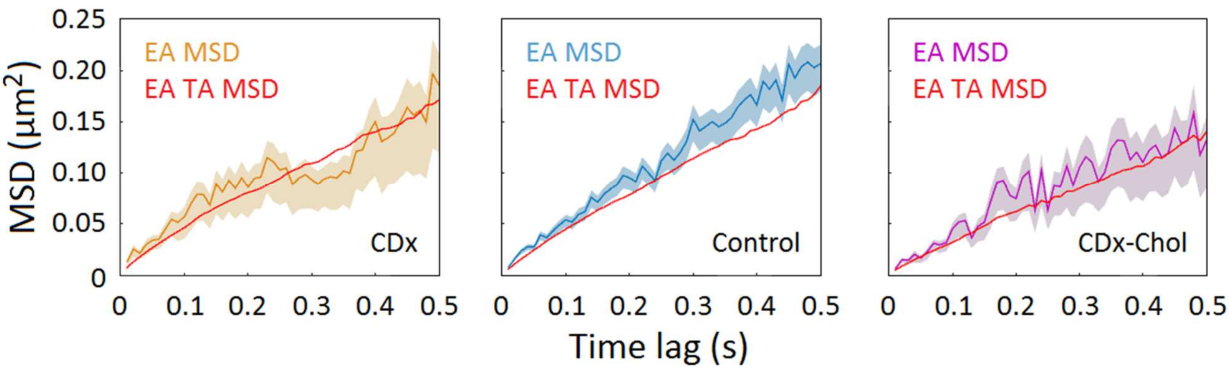


**Supplementary Figure 4. Mean-square displacement analysis applied to single-molecule nAChRs under control and cholesterol-modifying conditions.** The ensemble-averaged MSDs (EA MSD) with the 95% confidence interval is shown in color. An additional ensemble-average to the TA MSD (EA TA MSD, red trace) is performed to test the ergodic hypothesis (8-15 cells, 55 to 149 trajectories).

**Turning angles**

Turning angle analysis was recently employed to study the correlation of experimentally determined single-molecule steps in voltage-gated potassium channels Kv1.4 and Kv2.1 (Sadegh et al. 2017) compared to numerical simulations of fractional Brownian motion (fBM) and obstructed diffusion (OD) models. Experimental results corresponding to the control and cholesterol- depleted (CDx) / enriched (CDx-Chol) conditions for single-molecule nAChR trajectories are shown in Suppl. Figure 5.

Figure 2 in the main text shows that it is the slope of the turning angle in the confined portion that conforms with the obstructed diffusion model, and that the free portions of the trajectories exhibit a nearly flat Brownian-like behavior.

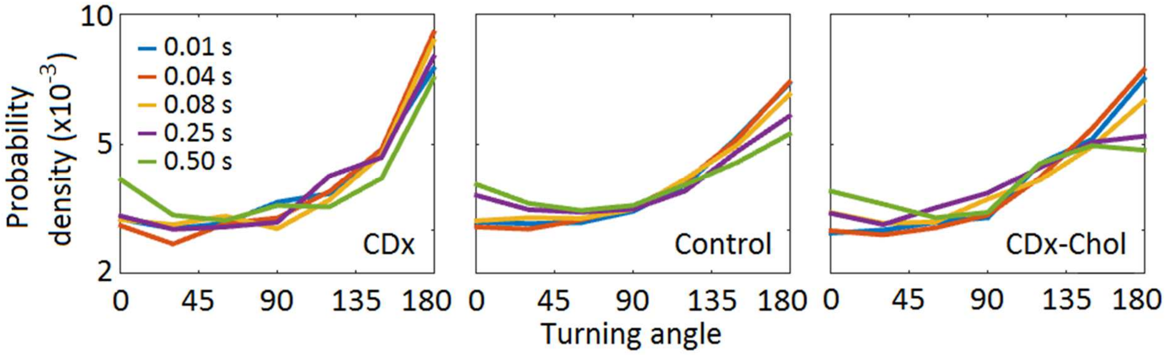


**Supplementary Figure 5. Turning angle probability densities of the single-molecule nAChR trajectories.** Probability densities for the color-coded $t_{lag}$ of increasing durations (10 to 500 ms) for the nAChRs under control and cholesterol modifying conditions (8-15 cells, 55 to 149 trajectories). The probability density is normalized such that the integral of the curve is equal to unity.
